# Computer-Assisted Engineering of Hyperstable Fibroblast Growth Factor 2

**DOI:** 10.1101/100636

**Authors:** Pavel Dvorak, David Bednar, Pavel Vanacek, Lukas Balek, Livia Eiselleova, Veronika Stepankova, Eva Sebestova, Michaela Kunova Bosakova, Zaneta Konecna, Stanislav Mazurenko, Antonin Kunka, Daniel Horak, Radka Chaloupkova, Jan Brezovsky, Pavel Krejci, Zbynek Prokop, Petr Dvorak, Jiri Damborsky

## Abstract

Fibroblast growth factors (FGFs) play numerous regulatory functions in complex organisms, and their corresponding therapeutic potential is of growing interest to academics and industrial researchers alike. However, applications of these proteins are limited due to their low stability *in vivo* and *in vitro*. Here we tackle this problem using a generalizable computer-assisted protein engineering strategy to create a unique modified FGF2 with nine mutations displaying unprecedented stability and uncompromised biological function.

## Introduction

The structurally related and highly conserved polypeptides from the family of FGFs are involved in a number of physiological processes in diverse animal species including humans. There has been a considerable effort in investigating members of the FGF family for applications in pharmacy and bioengineering^1^. Specifically, human FGF2 serves as a pleiotropic regulator of proliferation, differentiation, migration, and survival in a variety of cell types and has been studied as a promising agent in treatment of cardiovascular diseases^2^, cancer^3^ and mood disorders^4^. It has also been shown to have efficacy in ulcer, wound^5,6^ and epithelium healing^7^ and is being routinely used as an essential component of cultivation media for human embryonic stem cells^8^.

The long-term maintenance of growth factors in the tissue or media is desirable for protein therapies and stem cell culturing, but is hindered by low thermal stability of the molecules and their limited half-life^1,9,10^. Stability of FGFs was shown to be enhanced by co-administration with heparin^11^, conjugation to heparin-mimicking polymers^12^, encapsulation in microspheres^13^, or fusion with proteoglycan^14^. However, these strategies possess various limitations. They might significantly affect protein activity and are of prime concern from both safety and economic standpoints. Development of soluble, heparin-independent FGF analogues with improved stability is clearly needed for broader use of these interesting molecules^1^.

Protein engineering is a powerful approach for protein stabilization^15,16^. Stabilization by engineering has been previously applied to highly unstable FGF1 and resulted in improved mitogenic activity half-life by introducing stabilizing mutations at N and C terminus β-strand interactions of a β-barrel architecture.^17,18^ Triple^19^ and quintuple^20^ FGF2 mutants with improved stability and up to 10-fold prolonged activity in cell culture were recently prepared by aligning protein sequences of wild-type FGF2 with previously reported stabilized FGF1 mutants or by employing and combining individual stabilizing mutations published elsewhere. Nevertheless, the true potential of state-of-the-art protein engineering strategies has not yet been exploited in the design of stable FGFs. Here, we describe engineering of a unique nine-point mutant of low molecular weight isoform FGF2 with melting temperature (*T*_m_) increased by 19°C and *in vitro* functional half-life at 37°C improved from 10 hours to more than 20 days. This was achieved by following a computer-assisted engineering strategy combining energy-based and evolution-based analyses^21^ with focused directed evolution (Fig. 1). We demonstrate that the developed molecule holds a great promise both for *in vitro* and *in vivo* applications.

**Figure 1.**
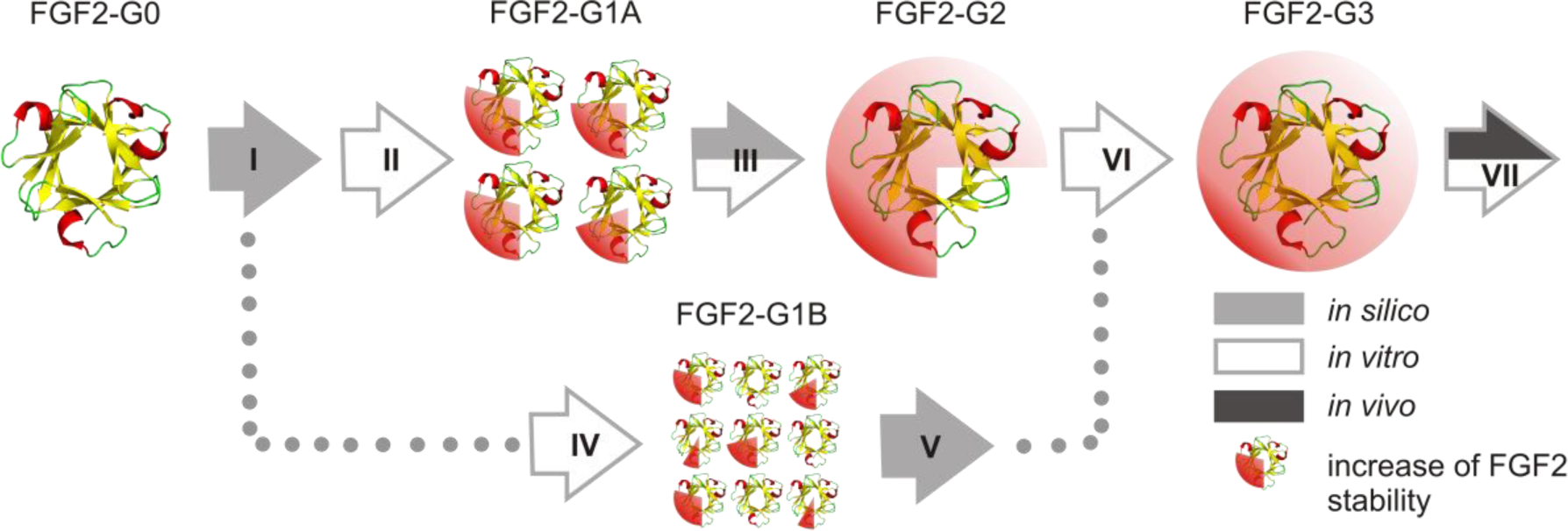
Integrated strategy combining computational analyses with focused directed evolution for engineering hyperstable FGF2. The workflow starts with the template wild-type molecule FGF2-G0. Initially, 12 computationally designed point mutants (FGF2-G1A) were constructed and tested leading to 7 stabilizing substitutions at 6 different positions with Δ*T*_m_ between 0.9 and 3.7°C. Six selected substitutions were recombined giving rise to the second generation mutant FGF2-G2 with Δ*T*_m_ of 15°C. Subsequently, 420 clones from 7 focused site-saturation mutagenesis libraries were screened for enhanced thermostability and retained biological activity (FGF2-G1B), revealing 14 beneficial substitutions at 5 randomized positions with *T*_m_ between 0.3 and 2.9°C. Guided by computational predictions, 5 substitutions from rational design were combined with 4 mutations from semi-rational design providing the third generation protein FGF2-G3 with Δ*T*_m_ of 19°C.

## Results and discussion

Using our platform, we proposed 12 single point mutations (R31L, R31W, V52T, H59F, C78Y, N80G, L92Y, C96Y, S109E, R118W, T121K, and V125L) and 7 positions for randomization (E54, C78, R90, S94, C96, T121, and S152) to stabilize FGF2 by computationally searching for mutations that would minimize the Gibbs free energy of the native state combined with a back-to-consensus analysis (Fig. 1 step I, Supplementary Figs. 1-2). Three out of twelve most stabilizing substitutions were identified by the phylogenetic approach^22^ and nine mutations were predicted from energetics using FoldX^23^ and Rosetta ddg monomer^24^ (Supplementary Tables 1 and 2). Seven positions with the highest number of potentially stabilizing mutations as defined by the difference in their free energy (ΔΔG) < -1 kcal.mol^-1^ were considered for randomization in order to look for further beneficial substitutions overlooked by the rational approach (Supplementary Table 2). Functionally relevant positions, namely those residues within heparin or receptor binding sites, were excluded from the designs to avoid mutations compromising activity.

For the purpose of *in vitro* mutagenesis (Fig. 1 steps II and IV), the gene encoding wild-type FGF2 was subcloned into vector pET28b with cleavable N-terminal His tag giving rise to the recombinant variant designated FGF2-G0 (Supplementary Fig. 2). The numbering of mutations used herein corresponds to the original sequence of wild-type FGF2 (Supplementary Fig. 2c). The genes of 12 rationally designed single-point mutants representing the first generation of engineered FGF2 (FGF2-G1A) were synthesized. Twelve mutants were produced in soluble form in *Escherichia coli* BL21(DE3) in quantities similar to that of FGF2-G0 (± 5% of the total soluble protein) and purified to homogeneity using affinity chromatography.

Biophysical characterization (Fig. 1 step II) verified proper folding of all 12 mutants and improved *T*_m_ for 7 out of 12 constructed variants (Supplementary Table 3). Six beneficial mutations (R31L, V52T, H59F, L92Y, C96Y, and S109E) were re-combined in the cumulative mutant FGF2-G2 (Fig. 1 step III), which was obtained at a yield of 20 mg.L^−1^. The experimental *T*_m_ of properly folded FGF2-G2 (68.0 ± 0.2°C) was about 14.5°C higher than *T*_m_ of FGF2-G0 (53.5 ± 0.5°C). This value corresponds well with the theoretical sum of contributions of individual substitutions, suggesting strictly additive stabilizing effect of mutations in FGF2-G2 (Supplementary Table 4).

In the next step, 7 computationally pre-selected positions in FGF2-G0 were randomized (Fig. 1 step IV) using fixed oligo technology, which reduced the screening effort needed to only 60 clones per library to obtain full coverage. Crude extracts of individual *E. coli* clones were prepared in microtiter plates and used to test simultaneously for biological activity and stability of FGF2 mutants in growth arrest assay with rat chondrosarcoma cells (Supplementary Fig. 3). Altogether, 33 new FGF2-G1B mutants were identified during the screening and successfully produced in *E. coli* to the levels similar to that of FGF2-G0. Thermal shift assays with purified proteins revealed 14 stabilizing amino acid substitutions in 5 out of 7 randomized positions (Supplementary Table 5). Mutations showing an improved thermostability of at least 1°C (E54D, S94I, C96N, and T121P) were computationally merged with existing mutations in FGF2-G2 (R31L, V52T, H59F, L92Y, C96Y, and S109E, while replacing C96Y by C96N) leading to the third generation variant FGF2-G3 (Fig. 1 steps V and VI and Fig. 2a).

**Figure 2.**
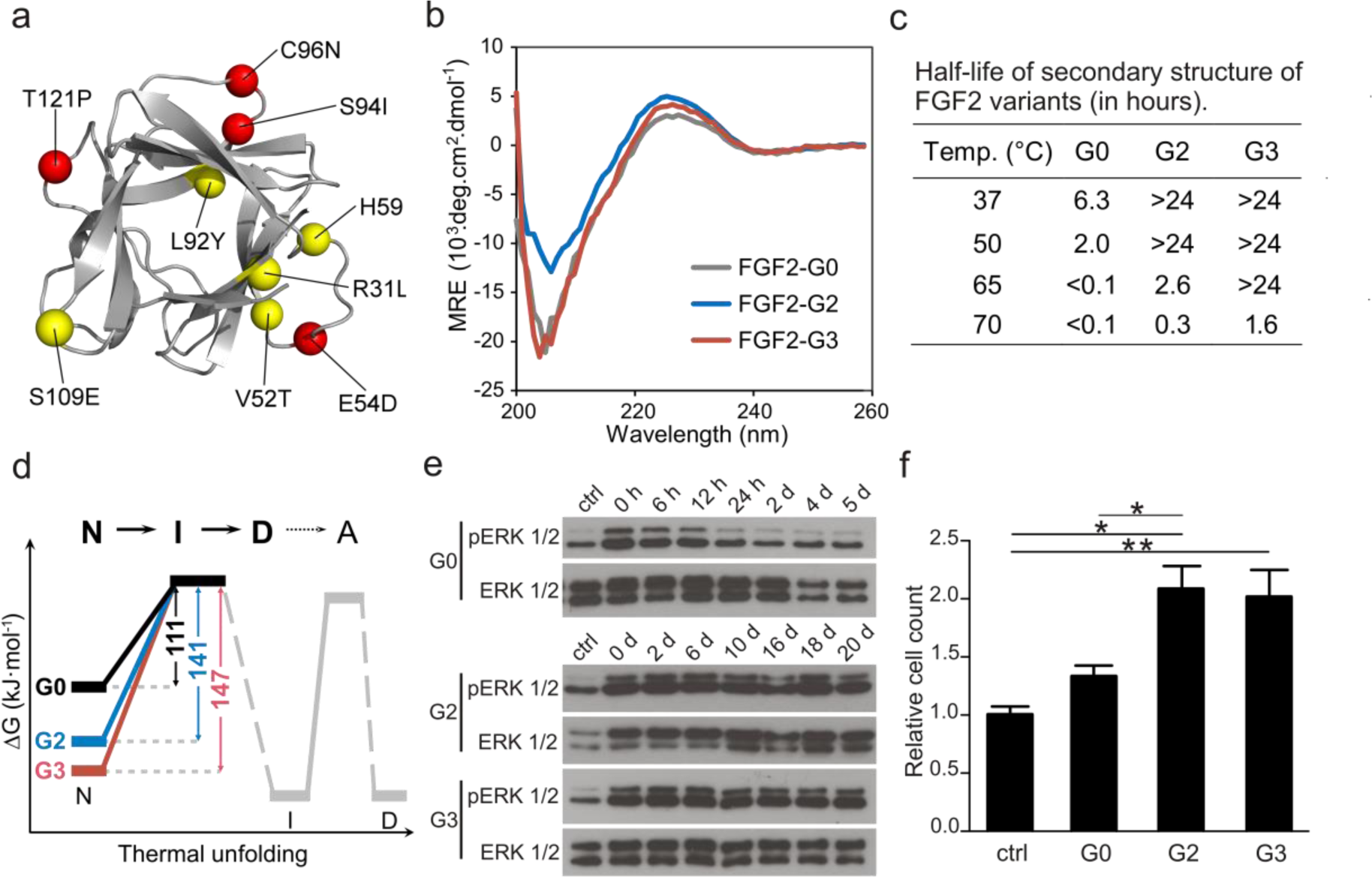
Biophysical and biological characteristics of FGF2 variants with boosted stability. (**a**) Structural model of the most stable FGF2-G3 variant with visualized amino acid substitutions originating from rational (yellow spheres) and semi-rational (red spheres) steps of the engineering strategy. (**b**) Circular dichroism (CD) spectra of selected FGF2 variants exhibiting a broad positive peak centered near 227 nm and a minimum at around 204 nm, characteristic for β-rich proteins of β-II type. (**c**) Half-life of FGF2 secondary structure of selected FGF2 variants determined by CD spectroscopy. (**d**) The schematic unfolding pathway of variants described by a three-step model with irreversible unfolding of native state (N) through one intermediate (I) to denatured protein (D) followed by formation of aggregates (A); Gibbs free energies of the states depicted in grey were not quantified because of irreversibility. (**e**) Determination of the *in vitro* functional half-life of selected FGF2 variants by activation of ERK pathway. Both FGF2-G2 and FGF2-G3 retained most of their original activity for 20 days of experiment, while FGF2-G0 lost half of its activity within initial 10 hours as determined by densitometric analysis of the corresponding Western blots (Supplementary Fig. 8). (**f**) FGF2-G2 and FGF2-G3 support proliferation of human embryonic stem cells better than the wild type. Columns show means, error bars represent standard error of the mean from three independent experiments. Student’s t-test, **p<0.01, *p<0.05. Abbreviations: MRE, mean residual ellipticity; ctrl, control with no FGF2; h, hours; d, days.

Purified FGF2-G3, obtained with the yield of 11 mg.L^-1^, was initially characterized *in vitro* using a wide spectrum of biophysical methods (Fig. 1 step VII). The properly folded mutant (Fig. 2b) exhibited a *T*_m_ of 72.2 ± 0.1°C and Δ*T*_m_ of 18.7°C compared with FGF2-G0, showing again the additive stabilizing effect of all new mutations, precisely predicted by the free energy calculation (theoretical Δ*T*_m_ of 19.2°C). In terms of structural integrity at elevated temperatures, FGF2-G3 clearly outperformed both FGF2-G0 and FGF2-G2, showing the half-life of its secondary structure of >24 h at 50°C and 1.6 h at 70°C (Fig. 2c and Supplementary Fig. 4). Fitting the data to a two-step unfolding mechanism suggested that gained stability was due to an increase in the Gibbs activation energy of the first unfolding step, which is in excellent agreement with our overall computational strategy of lowering the energy of the native state (Fig. 2d, Supplementary Results, Supplementary Table 6 and Figs 5-7).

Biological assays confirmed the enhanced stability of engineered growth factors translated into their prolonged activity *in vitro* and *in vivo* (Fig. 1 step VII). Specifically, while the activity of FGF2-G0 in an ERK1/2 assay with human embryonic stem cells dropped to 50 *%* within initial 10 ± 2 hours of pre-incubation in conditioned medium at 37°C (Fig 2e and Supplementary Fig. 8), which is in good agreement with data obtained previously by different techniques^1,25^, G2 and G3 mutants retained most of their activity for the full length of the study period (20 days) at the physiological temperature. In the second assay, the human embryonic stem cells were propagated in medium conditioned with each of the tested FGF2 variants with no additional supplementation, and the cell numbers and morphology were recorded for five consecutive passages (Fig. 2f and Supplementary Fig. 9). While conditioned medium prepared with FGF2-G0 caused significant growth retardation, the cells incubated in the medium with either G2 or G3 mutant gave rise to monolayers, suggesting that repeated supplementation of the conditioned medium is not required with these proteins. Immunostaining for Oct-4 and Nanog after five passages on Matrigel proved that all tested FGF2 variants support expression of pluripotency markers (Supplementary Fig. 10).

The biological activity of the stabilized variant was confirmed *in vivo* by injecting either FGF2-G0 or FGF2-G3, sorbed into a non-biodegradable hydrogel, into the shaved dorsal skin of telogenic C57BL/6 mice (Supplementary Fig. 11 and 12). It is known that growth factors from FGF family have positive effect on hair growth and hair follicle stimulation.^26^ We evaluated the degree of black pigmentation and hair growth photometrically by observing the skin color for 20 days. In 2 weeks, both FGF2 induced black coloration in the shaved skin with remarkably stronger hair growth induction in group injected with thermostable FGF2 (Fig. 3a, Supplementary Fig. 12). The control group with empty hydrogel showed significant retardation in transition from telogen to anagen stage as seen on skin coloration. The hair length of plucked hair in the proximity of injected site confirmed that thermostable FGF2 stimulated growth of hairs more significantly comparing to FGF2-G0 and the control group with no FGF2 applied (Fig. 3b). No toxic effects of recombinant proteins were observed.

**Figure 3.**
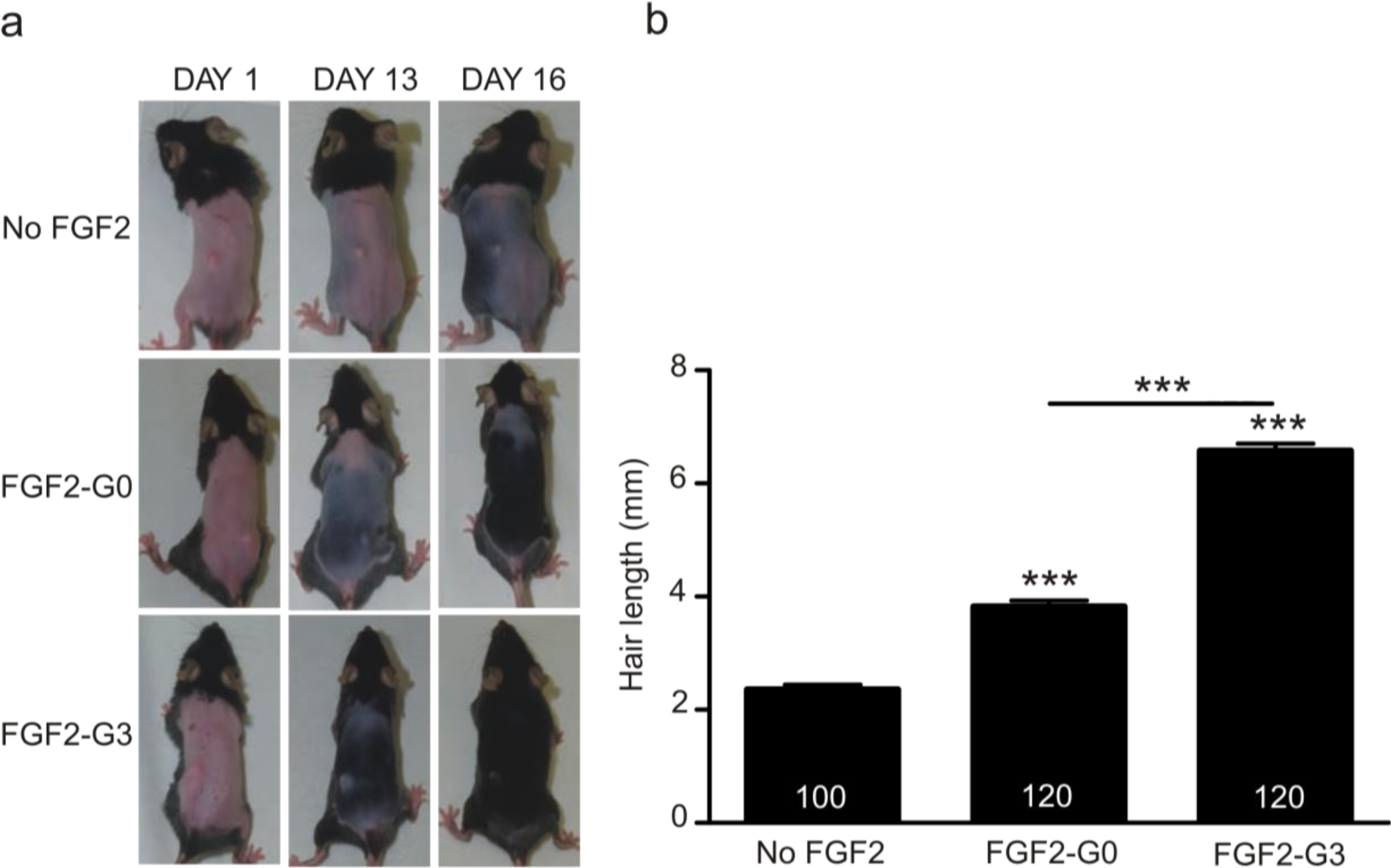
Effect of the selected FGF2 variants on hair growth promotion. **(a)** 7-week old C57BL/6 mice were shaved and injected with empty hydrogel or hydrogel sorbed with FGF2-G0 or FGF2-G3 and the hair growth was recorded during 20 consecutive days. **(b)** Hair length of C57BL/6 mice measured at the day 20 post application of FGF2 variants. Hairs (n=20) were plucked in the proximity of injection site. Columns show means, numbers within indicate the quantity of hairs counted per group, error bars show standard error of the mean. Student’s t-test,***p<0.001 versus control group.

In summary, we constructed a hyperstable nine-point mutant FGF2-G3 using a hybrid computational engineering strategy and focused directed evolution by constructing only 14 mutants and testing only 420 clones. The FGF2-G3 shows a 19°C increase in melting temperature and greater than 48-fold improved half-life at 37°C, representing the most thermostable FGF2 with fully preserved biological activity known to date. Our hyperstable FGF2 supports the undifferentiated growth of human embryonic stem cells, induces the appropriate downstream signaling cascade, and stimulates hair growth in mouse model, demonstrating that its biological activity towards highly sensitive cells remained unmodified. We anticipate this construct will be directly applicable to stem cell culturing^13^, and will find broad use in clinical medicine^1,12^, cosmetics^26^, and dietary supplements. The successful demonstration of rapid protein stabilization highlights the power of our rational protein engineering strategy and should encourage wider use of the described workflow for stabilizing growth factors and other protein therapeutics that are currently being tested for use in cancer treatment, regenerative medicine, or a number of metabolism-associated disorders^1^.

## Materials and methods

### Prediction of stabilizing effect of single-point mutations by evolution-based approach

#### Multiple sequence alignment and evolutionary conservation analysis

The FGF2 isoform 3 protein sequence (UniProt identifier: P09038-2) was used as a query for PSI-BLAST^28^ search against nr database of NCBI. PSI-BLAST was performed with the E-value thresholds of 10^−1^ for the initial BLAST search and the threshold of 10^−5^ for inclusion of the sequence in the position specific matrix. Sequences collected after 3 iterations of PSI-BLAST were clustered by CD-HIT^29^ at the 90% identity threshold. Resulting dataset of 554 sequences was clustered with CLANS^30^ using default parameters and varying P-value thresholds. Sequences clustered together with FGF2 at the P-value of 10^−30^ were extracted and aligned with the MUSCLE program^31^. The alignment was refined manually in BioEdit^32^. All incomplete or diverged sequences were removed. The final alignment comprising 238 sequences was used to estimate the level of conservation of individual sites within the FGF2 related proteins. Relative evolutionary rates for individual positions were calculated by the Rate4Site v2.01 program^33^ using the empirical Bayesian method^34^ and WAG model of evolution^35^. The evolutionary rates were then converted to the ConSurf conservation scale^36^.

#### Selection of individual mutations

The multiple sequence alignment comprising 238 FGF2 protein sequences was used as an input for back-to-consensus analysis using the simple consensus approach. The analysis was performed using the consensus cut-off of 0.5, meaning that a given residue must be present at a given position in at least 50% of all analysed sequences to be assigned as the consensus residue. Stability effects of all possible single-point mutations in FGF2 protein were estimated by free energy calculations (see following section for the calculation details). Only mutations with average Gibbs free energy (ΔΔG) ≤ 1 kcal.mol^−1^ predicted by both FoldX^22^ and Rosetta^23^ were considered as hot-spots for FGF2 stabilization. Functionally important sites of FGF2 were excluded as potentially deleterious mutations for biological function. Results of the back-to-consensus analysis are summarized in Supplementary Table S1. The numbering corresponds to the sequence of wild-type human FGF2 (Supplementary Fig. 2c). Ten mutations were excluded based on the high value of predicted ΔΔG or its high uncertainty of the prediction, and three mutations were discarded from the design due to their location at functionally important positions for the heparin binding. Four single-point mutations V52T, N80G, L92Y and S109E passed all criteria and were selected for experimental construction and characterization.

### Prediction of stabilizing effect of single-point mutations by energy-based approach, selection of positions for randomization

Available structures of FGF2 with resolution higher than 2.20 Å (PDB-ID codes: 1BFG, 4FGF, 2FGF, 1BAS, 1BFB, 1BFC, 1BFF, 1EV2, 1FGA) were downloaded from the RCSB Protein Data Bank^37^. The structures were visualized in PyMOL molecular graphics system v 1.7.7.4 (Schrödinger LLC, USA) and prepared for analyses by removing ligands and water molecules. Chain A was chosen in the case of multiple chain structure. Missing atoms in side chains were added by <RepairPDB> module of FoldX^22^.

#### Prediction of stability effects by FoldX

Stability effects of all possible single-point mutations were estimated using the FoldX <BuildModel> module. Calculations were performed 5-times for each mutant following the recommended protocol (pH 7, temperature 298K, ion strength 0.050 M, VdWDesign 2). For each mutation, the total ΔΔG value was calculated by averaging all ΔΔG values obtained for a respective mutation in all analyzed FGF2-G0 crystal structures.

#### Prediction of stability effects by Rosetta

Repaired structures were minimized by the minimize_with_cst module of Rosetta^23^ with both backbone and side-chains optimization enabled (--*sc_min_only false*), distance for full atom pair potential set to 9 Å (--*fa_max_dis 9.0*), standard weights for the score function and a constraint weight of 1 (--*constraint_weight 1.0*). Output from the minimization was used to constraint Cα atoms with harmonic function within 0.5 Å distance from the initial position in the crystal structure. Protocol 16 incorporating backbone flexibility within the ddg_monomer module of Rosetta was applied according to Kellogg and co-worker^23^. The soft-repulsive design energy function (--*soft_rep_design weights*) was used for repacking side-chains (--*sc_min_only false*). Optimization was performed on each whole protein without distance restriction (--*local_opt_only false*). The previously created constraint file was used during backbone minimization (--*min_cst true*). Three rounds of optimization with increasing weight on the repulsive term (--*ramp_repulsive true*) were applied. The minimum energies from 20 iterations were used as the final parameters describing the stability effects of single-point mutations.

#### Selection of individual mutations

Mutations predicted as stabilizing by either FoldX or Rosetta tool (ΔΔG < -1.0 kcal.mol^−1^) and not significantly constrained during evolution (Consurf conservation score < 8) were selected for further analysis. In this way, the potentially stabilizing mutations with only a limited influence on functional regions, *e.g.*, heparin binding residues, were identified. Residues forming the FGF2/FGFR1 interface (PDB-ID code 1CVS) and FGF2/FGFR2 interface (PDB-ID code 1EV2) were identified using the PISA server^38^ and were discarded from the selection. Nine single-point substitutions were selected for experimental construction and characterization: R31W, R31L, H59F, C78Y, L92Y, C96Y, R118W, T121K and V125L (Supplementary Table 2). Interestingly, the L92Y mutation was identified also previously by evolutionary-based approach. The numbering of these mutants corresponds to the sequence of wild-type human FGF2 (Supplementary Fig. 2c).

#### Selection of positions for saturation mutagenesis

Positions for saturation mutagenesis that should reveal additional stabilizing mutations were proposed using Rosetta calculations. Every protein position was saturated by all twenty proteinogenic amino acids and number of stabilizing mutations (ΔΔG < -1.0 kcal.mol^−1^) was identified for individual positions. Positions with conservations score > 7 or situated on the functional regions were discarded. Seven positions with the highest number (> 3) of stabilizing mutations (E54, C78, R90, S94, C96, T121, and S152) were selected for saturation mutagenesis (Supplementary Table 2). Positions 31 and 59 were discarded from selection because significant improvement in thermostability Δ*T*_m_=4°C and 3°C, respectively) was verified experimentally for the mutations R31L and H59F (Supplementary Table 3). Therefore, the probability of further considerable improvement was negligible.

### Construction, production and characterization of single point FGF2-G1A variants

Twelve FGF2-G1A variants R31W, R31L, V52T, H59F, C78Y, N80G, L92Y, C96Y, S109E, R118W, T121K and V125L were commercially synthesized (GeneArt/Life Technologies, Germany) and subcloned in the *NdeI* and *XhoI* sites of pET28b-His-thrombin downstream inducible T7 promotor. *E.coli* BL21(DE3) cells were transformed with expression vectors, plated on agar plates with kanamycin (50 μg.ml^−1^) and grown overnight at 37°C. Single colonies were used to inoculate 10 ml of LB medium with kanamycin and cells were grown overnight at 37°C. Overnight culture was used to inoculate 200 ml of LB medium with kanamycin. Cells were cultivated at 37°C. The expression was induced with IPTG to a final concentration of 0.25 mM. Cells were then cultivated overnight at 20°C. At the end of cultivation, biomass was harvested by centrifugation and washed by purification buffer A (20 mM di-potassium hydrogenphosphate and potassium dihydrogenphosphate, pH 7.5, 0,5 M NaCl, 10 mM imidazole).

Cells in suspension were disrupted by sonication using ultrasonic processor Hielscher UP200S (Teltow, Germany) with 0.3 s pulses and 85 % amplitude. Cell lysate was centrifuged for 1 h at 21,000 g at 4°C. FGF2 variants were purified from crude extracts using single step nickel affinity chromatography. Crude extracts were applied to a 5 ml Ni-NTA Superflow column (QIAGEN, USA). Column was attached to FPLC Akta (Amersham Pharmacia Biotech, USA). The buffer system consisted of buffer A and buffer B (20 mM di-potassium hydrogenphosphate and potassium dihydrogenphosphate, pH 7.5, 0,5 M NaCl, 500 mM imidazole). FGF2 proteins were eluted with a one-step increasing linear gradient of 0 to 100 % buffer B in 20 column volumes. The presence of FGF2 in peak fractions was proved by SDS-PAGE using 15 % polyacrylamide gel stained with Coomassie Brilliant Blue R-250 dye (Fluka, Buchs, Switzerland). Fractions with FGF2 were pooled and concentration of total protein was determined by Bradford method (Sigma-Aldrich, St. Louis, USA). Precipitation of FGF2 variants was minimized by dialysis against 20 mM potassium phosphate buffer containing 750 mM NaCl. Purified proteins were stored at 4°C.

#### Differential scanning calorimetry and circular dichroism spectroscopy

The thermostability of FGF2-G1 mutants was determined by differential scanning calorimetry (DSC) assay. Thermal unfolding of 1.0 mg.ml^−1^ protein solutions in 50 mM phosphate buffer (pH 7.5) with 750 mM sodium chloride was followed by monitoring the heat capacity using the VP-capillary DSC system (GE Healthcare, USA). The measurements were performed at the temperatures from 20 to 80°C at 1°C.min^−1^ heating rate. *T_m_* was evaluated as the top of the Gaussian curve after manual setting of the baseline. Proper folding of all mutants was verified by circular dichroism (CD) spectroscopy. CD spectra of mutants dialyzed in 50 mM phosphate buffer pH 7.5 and diluted to the concentration of 0.2 mg.ml^−1^ were recorded at 20°C using a spectropolarimeter Chirascan (Applied Photophysics, United Kingdom) equipped with a Peltier thermostat. Data were collected from 200 to 260 nm, at 100 nm.min^−1^, 1 s response time and 2 nm bandwidth using a 0.1 cm-quartz cuvette. Each spectrum is the average of five individual scans and is corrected for absorbance caused by the buffer. Collected CD data were expressed in terms of the mean residue ellipticity. Thermal unfolding was followed by monitoring the ellipticity at 232 nm over the temperature range from 20 to 80°C at a heating rate 1°C.min^−1^. Recorded thermal denaturation curves of FGF2 variants were normalized to represent signal changes between approximately 0 and 1 and fitted to sigmoidal curves. The melting temperatures were evaluated from the collected data as a midpoint of the normalized thermal transition.

### Free energy calculations, construction and thermostability analysis of FGF2-G2 mutant

All mutations improving the melting temperature by at least 0.5°C were selected for *in silico* analysis using Rosetta ddg monomer application^23^ as described earlier. In the case of two stabilizing mutations on the same position, the mutation with the larger effect was chosen. Selected mutations were then tested for potential antagonistic effect. The additivity of stabilizing mutations was evaluated by predicting the stability of variants with all pairs of stabilizing single-point mutations (Supplementary Table 4). Mutation pairs for which the respective double-point mutants showed lower stability in the comparison with the sum of both single-point mutants taken separately would be considered as antagonistic. However, none of the double-point mutants had the difference > 1 kcal.mol^−1^ suggesting the absence of any significant antagonistic effects among the selected mutations. All mutations improving *T_m_* by at least 0.5°C (R31L, V52T, H59F, L92Y, C96Y and S109E) were combined into 6-point mutant designated FGF2-G2. The gene of multiple-point mutant was commercially synthesized (GeneArt/Life Technologies, Germany), subcloned in the *NdeI* and *XhoI* sites of pET28b-His-Thrombin and expressed in *E. coli* BL21(DE3) cells as described before. DSC was used to characterize protein thermal stability of protein purified by affinity chromatography. DSC data collection was performed over a temperature range of 20°C–100°C.

### Construction and screening of focused site-saturation mutagenesis libraries

Altogether 7 focused site-saturation mutagenesis libraries were constructed commercially using “Fixed Oligo” technology of GeneArt (Life Technologies, USA). The pET28b-His-thrombin:*fgf2* was used as a template for randomization. Plasmid DNA was transformed into *E. coli* XJb (DE3) Autolysis cells (Zymo Research, USA). Cells were streaked on LB agar plates with kanamycin (50 μg.ml^−1^) and incubated overnight at 37°C. Plates with colonies carrying negative control (empty pET28b), positive control (plasmid pET28b-His-thrombin:*fgf2-G2*) and background control (pET28b-His-thrombin:*fgf2*) were prepared correspondingly.

#### Preparation of libraries for screening

Single colonies were used for inoculation of individual wells in 1 ml 96 deep-well plates (Thermo Fisher Scientific, USA) containing 250 μl of LB medium with kanamycin (50 μg.ml^−1^). Colonies were transferred by colony picking robot CP7200 (Hudson Robotics, USA) or using sterile wooden toothpicks. Plates were incubated overnight at 37°C with shaking (200 rpm) in shaking incubator NB-205 (N-Biotec, South Korea) in high humidity chamber to avoid evaporation of the medium. After 16 hrs, 50 μl of culture from each well was transferred to the new microtiter plate containing 50 μl of sterile 40 % glycerol per well and resulting plates were stored at -70°C as replicas. Expression of chromosomally inserted λ lysozyme and mutant variants of FGF2 in original plates with remaining 200 μl of overnight culture was induced by addition of 800 μl of fresh LB medium with kanamycin, IPTG and L-arabinose to the final concentration of 50 μg.ml^−1^, 0.25 mM and 3 mM, respectively. Plates were incubated overnight at 20°C with shaking (180 rpm). After 22 hrs, the plates were centrifuged for 20 min (3000 g, 4°C) using Sigma 6-16K (Sigma Laborzentrifugen, Germany). Supernatant was drained using JP recirculating water aspirator (VELP Scientifica, Italy). Whole microtiter plates with cell pellets were frozen at -70°C. Then, plates were incubated for 20 min at room temperature and 100 μl of lysis buffer (20 mM sodium phosphate buffer, 150 mM NaCl, pH 7.0) was added into the each well. Plates were incubated for 20 min at 30°C with shaking (200 rpm). Cell debris was removed from resulting cell lysates. Concentration of total soluble protein in crude extract in one well with negative control, one well with positive control, one well with background control and in 6 randomly selected wells with new FGF2-G1B mutants was determined for each plate containing one of the libraries using Bradford reagent (Sigma Aldrich, USA). Samples of crude extracts from the same wells were loaded on SDS polyacrylamide gels. The gels were analysed using GS-800 Calibrated Densitometer (Bio-Rad, USA) and the content of FGF2 in the total soluble protein in each sample was determined. The concentration of FGF2 in crude extracts was calculated based on the obtained data. The plates with crude extracts were stored at -70°C for further use.

#### Screening of biological activity of FGF2-G1B variants using rat chondrosarcoma growth-arrest assay

Rat chondrosarcoma (RCS) cells is an immortalized phenotypically stable cell line that responds to minute concentrations of FGFs with potent growth arrest accompanied by marked morphological changes and extracellular matrix degradation. FGF receptor 3 (FGFR3) functions as a negative regulator of cell proliferation in this cell line. In order to inhibit cell proliferation, FGF variants have to specifically induce FGFR signal transduction allowing the measuring of FGF activity reflected by the concentration dependence of induced growth arrest. The major advantage of the RCS assay is the exclusion of toxic chemicals and false-positive hits^39^. The high-throughput growth arrest experiment was performed in a 96-well plate format with the cellular content determined by simple crystal violet staining. Media with or without bacterial crude extracts with variants of FGF2 in approximate concentration of 40 ng.ml^-1^ were incubated at 41.5 °C for 48 h and mixed every 12 h within this period. RCS cells were seeded in concentration 250 cells per well in 96-well plate, one day before the treatment. Cells were treated with preincubated FGF2 at a final concentration of 20 ng.ml^-1^ for 4 days. Cells were washed with PBS, fixed with 4% paraformaldehyde, washed again and stained with 0.025% crystal violet for 1 hour. Coloured cells were 3 times washed with distilled water. Colour from cells was dissolved in 33% acetic acid. Absorbance was measured at 570 nm (Supplementary Fig. 3). The more stable variant of FGF2 was present in added crude extract, the more evident was the growth inhibition. Samples causing more significant growth inhibition than samples containing FGF2-G0 were considered as positive hits. *E. coli* clones containing FGF2 candidates were refreshed from glycerol replica plates. For each of positive hits, four wells with LB medium in fresh 1 ml 96-well PP microtiter plate were inoculated and the whole screening procedure was repeated. For each of FGF2 candidates verified in second round of screening, 10 ml of LB medium with kanamycin was inoculated with corresponding *E. coli* clone from glycerol replica plate and the cells were grown overnight at 37°C with shaking. The overnight culture was used for isolation of plasmid DNA using GeneJET Plasmid Miniprep kit (Thermo Fisher Scientific, USA) and *fgf2-G1B* genes were commercially sequenced by Sanger method (GATC Biotech, Germany). Resulting sequences were aligned with nucleotide sequence of FGF2-G0 using BioEdit32 for determination of newly inserted mutations.

### Small scale production and characterization of selected FGF2-G1B mutants

*E. coli* BL21(DE3) cells were transformed with pET28b-His-thrombin:*fgf2x* (where x represents one of 33 new mutant variants), plated on LB agar plates with kanamycin (50 μg.ml^−1^) and grown overnight at 37°C. Small scale cell cultivations in 10 ml of LB medium with kanamycin was conducted under conditions described before. The biomass was centrifuged at 10,000 g for 2 minutes at 4°C in a benchtop centrifuge Mikro 200 (Andreas Hettich GmbH & Co.KG, Germany) and the cell pellet was frozen at −70°C. The pellets were defrosted and resuspended in 600 μl of FastBreak Cell Lysis Reagent from MagneHis Protein Purification System (Promega, USA) added with NaCl to the concentration of 500 mM and 1μl of DNase I (New England Biolabs, USA). The cells were incubated with shaking for 20 minutes at room temperature. The bacterial lysates were incubated with 30 μl of MagneHis Ni-Particles beads for 2 minutes at room temperature. The beads were separated using magnetic stand and the supernatants were carefully removed. To wash out unbound cell proteins, 150 μl of MagneHis Binding/Wash Buffer with 500 mM NaCl was added. The elution of bound proteins was performed by adding 105 μl of MagneHis Elution Buffer containing 500 mM NaCl. The presence of FGF2-G1B variants in eluted fractions was proved by SDS-PAGE as described before.

#### Determination of thermal stability of FGF2-G1B mutants

The thermal stability of FGF2-G1B variants was verified by thermal shift assay^40^. FGF2-G0 was used as a background control. The measurements were conducted in MicroAmp Fast Optical 96-well Reaction Plate (Thermo Fisher Scientific, USA). Each reaction mixture of final volume of 25 μl was composed of 2 μl of SYPRO Orange Protein Gel Stain (Thermo Fisher Scientific, USA), purified FGF2 variant (2.5 mg.ml^−1^) and the elution buffer (100 mM HEPES, 500 mM imidazole and 500 mM NaCl, pH 7.5). The assay was performed using StepOnePlus Real-Time PCR System (Applied Biosystems/Thermo Fisher Scientific, USA) with starting temperature of 25°C (2 min initial equilibration) and ramping up in increments of 1°C to a final temperature of 95°C. The *T*_m_ values were determined from obtained data using Protein Thermal Shift software (Applied Biosystems/Thermo Fisher Scientific, USA; Supplementary Table 5).

### Free energy calculations, construction and purification of FGF2-G3 mutant

All mutations from screening improving the melting temperature by at least 1°C (E54D, S94I, C96N, and T121P) were selected for *in silico* analysis using Rosetta ddg monomer application^23^ as described earlier. All double-point mutant combinations of newly identified mutations with existing mutations from stable variant FGF2-G2 (R31L, V52T, H59F, L92Y, C96Y, and S109E) were constructed *in silico* to predict potential additivity of these individual mutations (data not shown). Predicted ΔΔG was compared with the sum of ΔΔG of individual mutations but none of the double point mutants had the difference > 1 kcal.mol^−1^ again suggesting the absence of any antagonistic effects among selected mutations. Consequently, 9-point mutant FGF2-G3 was designed and constructed combining 4 most stabilizing substitutions obtained from screening with 5 substitutions from FGF2-G2. In FGF2-G3 mutant, the substitution C96N was prioritized over C96Y due to its higher individual stabilizing effect verified experimentally. Predicted improvement in *T*_m_ for this new variant was 19.2°C. The gene of multiple-point mutant was commercially synthesized (GeneArt/Life Technologies, Germany), subcloned in the *NdeI* and *XhoI* sites of pET28b-His-Thrombin and expressed in *E. coli* BL21(DE3) cells as described before.

### Characterization of biophysical properties of FGF2-G3 and its comparison with FGF2-G0 and FGF-G2 variants

DSC was used to characterize protein thermal stability of protein purified by affinity chromatography. DSC data collection was performed over a temperature range of 20°C-100°C. Proper folding of mutant was verified by CD spectroscopy as described earlier.

#### Circular dichroism spectroscopy

The structural integrity of FGF2-G0, FGF2-G2 and FGF2-G3 proteins was followed by monitoring the ellipticity over the wavelength range of 200 to 260 nm at the temperature 37, 50, 65 and 70°C for 24 h. Data were recorded in 2 minute intervals with 1 nm bandwidth using a 0.1 cm quartz cuvette containing the protein. Recorded denaturation curves (either single exponential, double exponential or exponential linear combination function) of tested FGF2 variants were globally fitted to exponential decay curves using OriginPro8 software (OriginLab, USA). Half-life of FGF2 secondary structure (*t*_1/2_ defined as a time required to reduce the initial value of ellipticity, as a measure of protein secondary structure, to ½ of the original value) was evaluated from the collected data as a decay constant (τ) using the equation:

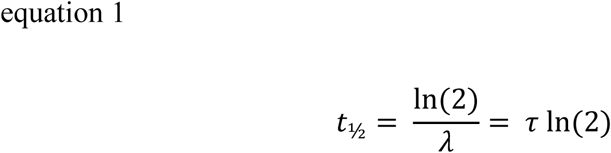

where λ is exponential decay constant. The *t*_½_ was evaluated from the data collected at 227 nm, where all spectra showed the ellipticity maxima (Supplementary Fig. 4). The maximum time of measurement of 24 h was limited by the capacity of the bomb with compressed nitrogen used in the CD spectroscopy protocol. The values of *t*_½_ for FGF2-G0 at 65 and 70 °C were not determined because the proteins were denatured immediately at the beginning of the measurement.

#### Fluorescence spectroscopy

Local conformational changes during thermal unfolding of FGF2 variants were followed by monitoring fluorescence emission spectra using FluoroMax spectrofluorometer (Horiba, Japan). The sample in quartz cuvette with a magnetic stirrer inside was placed into a temperature-controlled holder, and fluorescence spectra excited at 295 nm were recorded in 1 minute intervals from 310 to 410 nm with 1 nm bandwith and 0.1 s integration time from 30 to 90 °C. The actual temperature in the cell was monitored using thermocouple controlled by Labview software (National Instruments, USA). The spectrum of the buffer recorded at 30 °C was used as a blank and subsequently subtracted from the sample data. Unfolding was followed at the emission maximum (347 nm) of the first scan. The concentration of all the samples was approx. 0.1 mg.ml^−1^.

#### Differential scanning fluorescence (DSF)

A slightly different experimental set-up was used for monitoring fluorescence during thermal unfolding. The standard grade capillary (NanoTemper, Germany) was filled with a sample and placed into the Prometheus NT.48 (NanoTemper, Germany). The samples were continually heated from 30 to 90 °C at different scan rates (0.3, 0.5, 1, 2 and 4 °C) and fluorescence signal excited at 295 nm was followed at 335 and 350 nm. The concentration dependence of the unfolding curve was checked by monitoring aliquots of different concentrations (1, 0.5, 0.25 and 0.125 mg.ml^-1^).

#### Data analysis

The data were uploaded to an extension of CalFitter (Masaryk University, Czech Republic) based on MATLAB 2014b (The MathWorks, United States) that allows simultaneous global fit into unfolding curves. DSC data were numerically integrated to derive the total heat absorbed during the transition. Then the signals from the four types of measurement, namely CD ellipticity, DSC heat absorption, and two fluorescence measurements, were normalized and fit globally (Supplementary Figs. 5 and 6). After the initial fitting, the weighted least squares were calculated based on the sum of the squared residuals of each curve; thereby, the contributions of each type of the measurements to the sum of errors were the same at the optimal point. Regarding the parameters, linear coefficients were allocated separately to each data set. The models with the minimum number of intermediates were selected based on the quality of fit measured by normalized residuals and visual inspection. Although the datasets of the wild type and the G2 mutant were fitted reasonably well with just a three-step irreversible model, the G3 model had to include one additional reversible pretransitional step to fit all the data sets perfectly, mainly due to the DSC data and ratiometric data from DSF. Nonetheless, the effect of this step is much less pronounced than the subsequent irreversible steps of unfolding in all the datasets. The estimated values of the main parameters of the unfolding mechanism are given in Supplementary Table 6.

#### pH profile

Britton-Robinson buffers of different pHs (5, 6, 7, 7.5, 8, 9, 10 and 11) were used for determination of pH stability profile for the FGF2 variants. 5 μL sample aliquot was mixed with 95 μL of BR buffer of appropriate pH, thoroughly vortexed and incubated for 3 h at 4 °C. Next standard grade capillary was filled with a mixture by capillary forces and put into the Prometheus NT.48. Samples were scanned from 30 to 90 °C at 1°C.min^−1^ scan rate.

### Characterization of biological properties of FGF2-G3 and its comparison with G0 and G2 variants

The authors herein declare that all following experiments conducted on animals or human cells were carried out in accordance with relevant guidelines and regulations. All experimental protocols were approved by Faculty of Medicine of Masaryk University and licensing committee of Czech Ministry of Health.

#### Determination of biological activity half-life

FGF-receptors and their downstream effectors including ERK1/2 are activated upon treatment with FGF2, contributing to pluripotency of human embryonic stem cells (hESC)^41,42^. As the biological activity of FGF2 decreases at 37°C, ERK1/2 phosphorylation declines and hESC easily become primed to differentiation. To test the thermal stability of FGF2 variants, the hESC medium prepared without FGF2 was supplemented with FGF2-G0, FGF2-G2 or FGF2-G3 to the final concentration of 10 ng.ml^−1^ and pre-incubated at 37°C for 6 h, 12 h, 24 h, 2 d, 4 d – 20 d. FGF2-starved hESC were treated with hESC medium containing pre-incubated FGF2 for two hours and Western blotted for phosphorylated ERK1/2. Two representative blots per each protein variant were analysed using ImageJ 1.50b (National Institutes of Health, USA) and band densities were plotted as a function of pre-incubation time (Supplementary Fig. 8). Data points were analysed using single exponential decay equation:

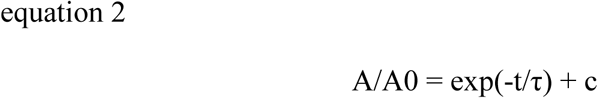

where A/A0 is the relative density at time t, τ is the time constant, and c is steady state level of density offset, with help of OriginPro8 software (OriginLab, USA) and the half-life (i.e. the time required for the loss of one-half of the initial activity) was determined for FGF2-G0 protein variant.

#### Western blotting

Cells were lysed with 2x Laemmli buffer and the samples were boiled at 98°C for 10 minutes. Proteins were separated by SDS-PAGE and electrotransferred onto Immobilon®-P transfer membrane (Merck Millipore, Germany). Membranes were then blocked in 5% milk in TBS buffer and incubated with primary rabbit polyclonal antibody P- 44/42 MAPK (Cell Signaling Technologies, USA) at 4°C overnight. The primary antibodies included rabbit polyclonal anti-pERK1/2 and rabbit polyclonal anti-ERK1/2 (both Cell Signaling Technology, USA). Next day, the membranes were incubated with donkey antirabbit antibodies conjugated with horse raddish peroxidase (Santa Cruz Biotechnology, USA), and the protein bands were visualized using chemiluminiscence detection reagent ImmobilonTM Western (Merck Millipore, Germany) on photographic paper (Agfa-Gevaert, Belgium). After stripping, the membrane was re-probed with rabbit polyclonal antibody p44/42 MAPK (Erk1/2) (Cell Signaling Technology, USA) against total signaling proteins.

#### Cell cultures

The hESC employed in this study were derived from blastocyst-stage embryos obtained with informed consent of donors. A well characterized human ESC line (Adewumi, 2007) CCTL14 (Centre of Cell Therapy Line) in passages 65 - 75 was used. The hESC were maintained under feeder-free conditions using MatrigelTM hESC-qualified Matrix (BD Biosciences). Culture medium required for propagation of hESC grown on Matrigel was medium conditioned by mitotically inactivated mouse embryonic fibroblasts (mEF). For preparation of standard conditioned medium (CM), the complete hESC medium containing 4 ng.ml^−1^ FGF2 is usually conditioned by mitotically inactivated mEF for 5-7 days and then supplemented by 10 ng.ml^−1^ of FGF2 to restore growth factor concentration due to its degradation. In our experiments, to test the long-term thermostability of FGF2, the CM was prepared out of medium containing 10 ng.ml^−1^ of FGF2 with no supplementation afterwards.

#### Proliferation assays

The hESC were plated into 24-well plates and propagated in presence of each of the tested FGF2 for five passages and counted every three days after plating using Burker chamber. Alternatively, cells were plated into 94-well plates and cultured in the presence of various FGF2 for 6 days. Cells were then fixed in 4% paraformaldehyde (20 min, RT), stained with 0.1% crystal violet (60 min, RT), and destained with 33% acetic acid (20 min with shaking). The absorbance of the supernatant was then measured at 570 nm using plate reader (Supplementary Fig. 9).

#### Immunocytochemistry

The hESC were fixed with 4% paraformaldehyde (20 min, RT), permeabilized with 0.1% Triton-X100 in PBS (20 min, RT), and incubated with primary antibodies at 4°C overnight. Primary antibodies included goat polyclonal anti-Oct4 (Santa Cruz Biotechnology, USA) and rabbit monoclonal anti-Nanog (Cell Signaling technology, USA). Next day, incubations with secondary antibodies conjugated to AlexaFluor488 or AlexaFluor594 (Thermo Fisher Scientific, USA) were carried out at RT for 1 h. Coverslips were mounted in DAPI-containing Mowiol (Sigma-Aldrich, USA). Microscopic analysis was performed using Confocal LSM 700 microscope (Zeiss, Germany; Supplementary Fig. 10).

#### Preparation of hydrogel

Macroporous poly(2-hydroxyethyl methacrylate) (PHEMA) microspheres of narrow particle size distribution were prepared by multi-step swelling polymerization.^43^ Briefly, the method is based on 0.7 μm monodisperse polystyrene seeds which are swollen with activating agent (dibutyl phthalate), monomers (2- (methacryloyl)oxyethyl acetate, 2-[(methoxycarbonyl)methoxy]ethyl methacrylate, ethylene dimethacrylate), and porogen (cyclohexyl acetate). After benzoyl peroxide-initiated and (hydroxypropyl)methyl cellulose-stabilized polymerization, hydrolysis, and washing, the resulting PHEMA microspheres were 3 μm in diameter, with a narrow size distribution (Supplementary Fig. 11) and contained 0.5 mmol COOH/g.

In vivo *experiment*. Animals used in the study of hair promoting activity, 7 weeks old female C57BL/6 mice, were obtained from Laboratory Animal Breeding and Experimental Facility (Masaryk University, Brno, Czech Republic) and maintained on a standard laboratory diet and water as libitum. 17 animals in 3 randomized groups (n=5 or 6) were shaved using depilatory cream (Veet, USA) at 7 weeks of age, at which all hair follicles were synchronized in the quiescence telogen.^44^ FGF2, both FGF2-G0 and thermostable variant FGF2-G3 (5 μg per mouse) were dissolved in 0,1% human serum albumin and sorbed into the hydrogel by continuous stirring for 2 hours at RT. The resulted suspension was applied topically on dorsal skin of C57BL/6 mice with subcutaneous injection. Empty hydrogel with no FGF2 was used as a control. Visible hair growth was recorded at days 1, 7, 13, 16 and 20 (Supplementary Fig. 12).

#### Hair length determination

To examine the effect of FGF2 in hydrogel on hair length, hairs on the proximity of injected site were plucked randomly by forceps at day 21 post application. The average length of the 20 plucked hairs per mouse was measured manually with a micrometer under a stereoscopic microscope and expressed in millimeters.

## ACKNOWLEDGEMENTS

We would like to thank Dr. Sergiy Kyrylenko (Masaryk University, Brno) for providing *pET28b::fgf2* construct and information on existing FGF2 mutants. The work was supported by the Grant Agency of the Czech Republic (GA16-06096S, GA15-23033S), Ministry of Education of the Czech Republic (LO1214, LQ1605, LM2015051, LM2015047, LM2015055), and Ministry of Health of the Czech Republic (15-33232A). MetaCentrum and CERIT-SC are acknowledged for providing computing facilities (LM2015042, LM2015085). The funders had no role in study design, data collection and analysis, decision to publish, or preparation of the manuscript.

## AUTHOR CONTRIBUTIONS

Conceived and designed the experiments: All authors. Performed the experiments or computational calculations: PaD, DB, PV, LB, LE, ES, MKB, ZK, SM, AK, DH and RC. Analyzed the data: All authors. Wrote the paper: PaD, DB, VS, SM, RC, JB, ZP, PK, PeD and JD.

## COMPETING INTERESTS

A patent application protecting intellectual property related to stabilized FGF2 has been submitted to European Patent Office on behalf of Masaryk University and Enantis Ltd. JD and ZP are founders and VS is an employee of the university spin-off company Enantis Ltd. All other authors have no competing interests.

